# AutoPELSA: an automated sample preparation system for proteome-wide identification of target proteins of diverse ligands

**DOI:** 10.64898/2026.01.08.697612

**Authors:** Lianji Xue, Xi Wang, Haiqian Yang, Jiahua Zhou, Haiyang Zhu, Jinhua Shen, Weina Gao, Ruijun Tian, Yan Wang, Mingliang Ye

**Author notes:** Corresponding authors: Yan Wang. Mingliang Ye.

## Abstract

Protein-ligand interactions are fundamental to cellular function and drug discovery, and ligand-modification-free strategies have emerged as powerful tools for proteome-wide interrogation of these interactions. Among these, the peptide-centric local stability assay (PELSA) stands out for its high sensitivity. It uses a single digestion step to detect ligand-induced local stability shifts, enabling precise binding region localization and affinity estimation. However, manual PELSA workflows suffer from multiple labor-intensive steps that introduce variability and constrain throughput, limiting large-scale applications. To address these limitations, we developed AutoPELSA, an automated platform that streamlines the PELSA workflow including the limited proteolysis step and the following peptide separation step. AutoPELSA completes 96 samples in ∼4 h, enabling high-throughput analysis under native conditions. AutoPELSA reliably detected targets of both strong-affinity ligands (e.g., staurosporine, identifying 114 kinase targets) and low-affinity ligands, including α-ketoglutarate (30 known targets). Furthermore, a mixed-ligand dose–response strategy enabled simultaneous determination of binding affinities for multiple ligand–protein interaction regions in a single experiment. Overall, AutoPELSA provides a scalable and modification-free platform for proteome-wide identification and affinity profiling of ligand–protein interactions.

## Introduction

Protein–ligand interactions are fundamental to cellular function, underpinning enzymatic catalysis, signaling, immunity, and gene regulation, and thus remain a central focus of chemical biology and proteomics^1–4^. Yet comprehensive target identification for diverse ligands remains a pressing need as resolving these targets directly informs drug design^5^, and a deeper understanding of protein–ligand binding facilitates the development of more effective therapeutics^6^ by enabling structure-guided optimization of affinity and selectivity, revealing off-target liabilities, and informing strategies to improve efficacy and safety^7^. With advances in MS-based proteomics and computational pipelines, ligand-modification–free strategies have become increasingly prevalent for large-scale interrogation of protein–ligand interactions^8^. Because ligand binding can induce local conformational and thermodynamic changes that modulate protease accessibility, limited proteolysis coupled to MS (LiP-MS)^9^ was developed to identify binding proteins and binding sites. However, LiP-MS relies on detecting specific peptide forms within complex peptide pool generated by the second digestion step, which compromises the sensitivity for confident target identification. In contrast, the peptide-centric local stability assay (PELSA) uses a single digestion step, i.e. extensive trypsinization of native proteins to directly generate MS-detectable peptides, to reveal the local stability changes induced by ligand binding^10^. This design enables localization of binding regions and estimation of local affinity, and has demonstrated much higher sensitivity than other ligand-modification-free methods for target identification, including weak interactions^11, 12^.

However, PELSA is constrained by multiple manual steps (lysis, ligand incubation, trypsin digestion, reduction/alkylation, ultrafiltration, and desalting), each step introduces variability. To partially mitigate this, researchers typically employ four technical replicates per condition to enhance measurement precision, reduce variance, and thereby boost statistical power. But this comes at the cost of low throughput: profiling one ligand requires 8 samples and ∼8 h of processing, limiting capacity to two ligands per day. For concentration–response designs (∼30 samples to quantify ligand-target binding affinity), the required multi-window processing introduces additional between-time-point variability. Further, precise control of the ∼1-min digestion window is particularly challenging with manual handling. These limitations motivate the development of high-throughput, standardized sample preparation for large-scale PELSA analysis.

Although many proteomic preparation workflows have been automated^13–15^, PELSA is not readily adaptable to these systems. Beyond the stringent digestion step, automation is hindered by the reliance on ultrafiltration-based protein removal, which is poorly suited for automated platforms. Researchers have explored single-cartridge C18 solid-phase extraction as a strategy to simultaneously remove proteinaceous interferents and desalt samples^16^, but C18 media alone are insufficient to eliminate large quantities of proteinaceous contaminants from complex digest mixtures.

To address these challenges, we developed AutoPELSA, an automated sample-preparation platform that integrates and streamlines the entire PELSA workflow. Redesigning the quenching strategy and optimizing working volumes, alleviated the barriers to automate the sub-minute digestion/quenches, thereby permitting reliable execution on a programmable liquid-handling platform and enhancing reproducibility. To further increase throughput, ultrafiltration and desalting were consolidated into a single operation using a custom C4–SCX–C18 multilayer tip. This architecture removes hydrophobic and highly charged proteinaceous fragments while simultaneously achieving peptide desalting. Consequently, AutoPELSA delivers a substantive throughput increase (∼20×), enabling completion of 96 PELSA samples within ∼4 h. Moreover, the multilayer tip design substantially lowers consumable costs, supporting cost-effective, large-scale ligand screening.

## Results and discussion

### Design of AutoPELSA

To improve the throughput and reproducibility of PELSA sample preparation, we developed AutoPELSA, which consists of two functionally integrated modules (Fig. 1a). The first module automates the limited proteolysis step on a liquid handling platform equipped with a 96-channel pipette, encompassing ligand incubation with cell lysates, precisely timed limited trypsinization, protein denaturation, reduction and alkylation, and final acidification. Accordingly, we systematically optimized the critical steps and adjusted the buffer volumes for sample preparation step to suit automated operation. The second module implements high-throughput cleanup with a tip-column format on a positive-pressure manifold. A multilayer tip design enables concurrent removal of partially digested proteins and residual trypsin, together with peptide desalting, in one streamlined operation and therefore allows the specific separation of the peptides from the complex digests for bottom-up proteomics analysis (Fig. 1b).

**Figure 1.**
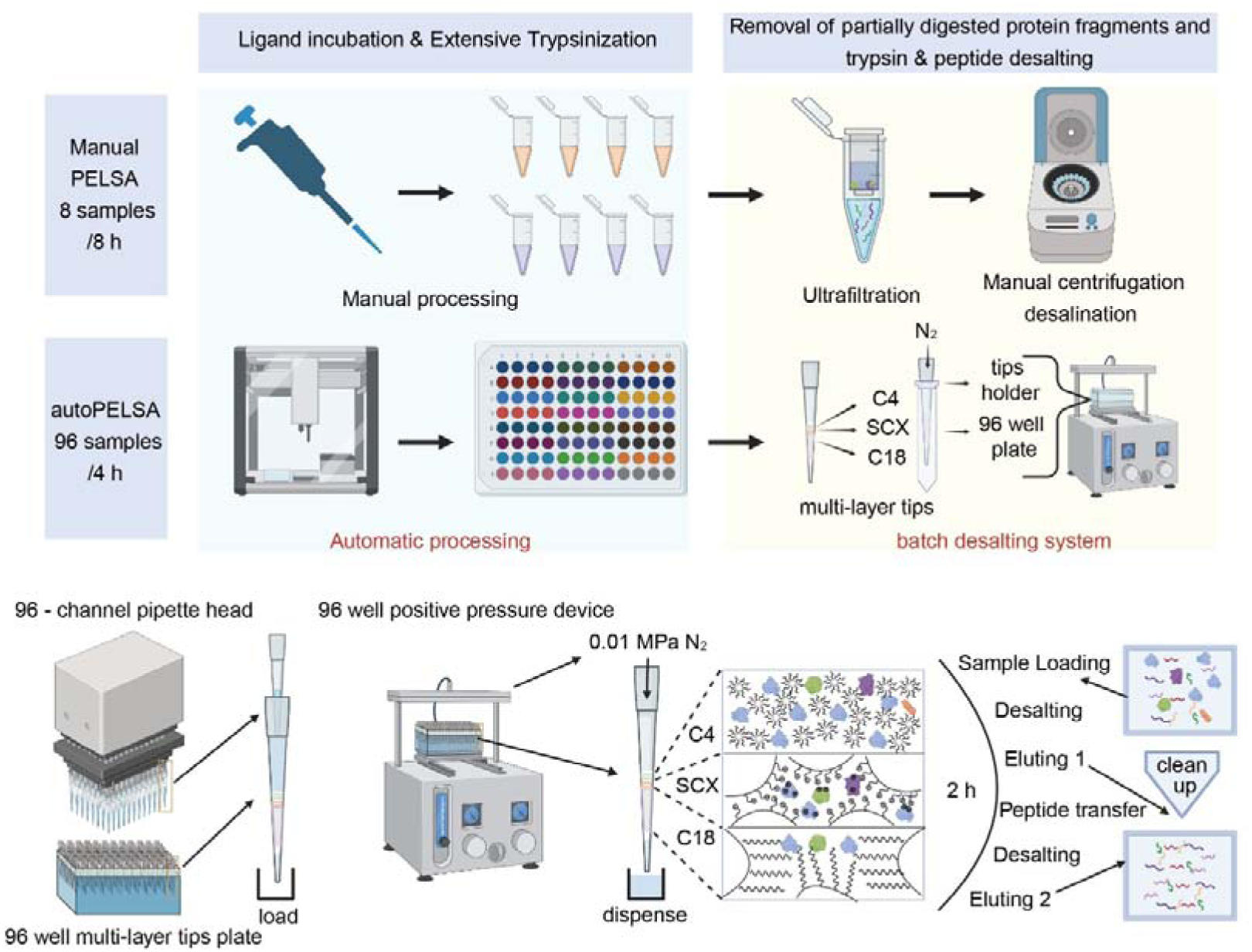
Overview of the PELSA sample preparation procedures and AutoPELSA workflows. (a) Comparison of manual PELSA and AutoPELSA. The manual workflow involves sequential pipetting, ultrafiltration, and centrifuge-based desalting. In AutoPELSA, ligand incubation and extensive trypsinization are executed on a 96-channel liquid-handling workstation, followed by cleanup using a C4–SCX–C18 multilayer tip operated in batch mode. (b) Design and operation of the multilayer tip under positive pressure. The stacked C4, SCX, and C18 layers sequentially retain proteinaceous fragments and enable peptide desalting through loading, washing, and two-step elution.

### Automation of the limited trypsinization on a Liquid-Handling Platform

Precise control of digestion time is critical for PELSA, so rapid and complete quenching of trypsin is essential. In the manual workflow, quenching is achieved by adding a large volume of guanidine hydrochloride to denature the trypsin. However, dispensing large volumes in a single step is not feasible on a liquid-handling workstation and would substantially compromise timing precision.

Alternatively, acid quenching with HCl is possible: trypsin is fully inhibited at pH ≤3^17^. In our workflow, adding a small volume of HCl (approximately 10 µL of 0.1 M) rapidly lowers the pH to this range and is compatible with standard pipetting workstations. To validate this approach, we compared the acid-quenching strategy with the conventional guanidine hydrochloride method under otherwise identical manual PELSA conditions. When profiling targets of the pan-kinase inhibitor staurosporine, the two methods exhibited equivalent performance, with each identifying about 90 kinase targets under identical experimental conditions (Fig. 2a).

**Figure 2.**
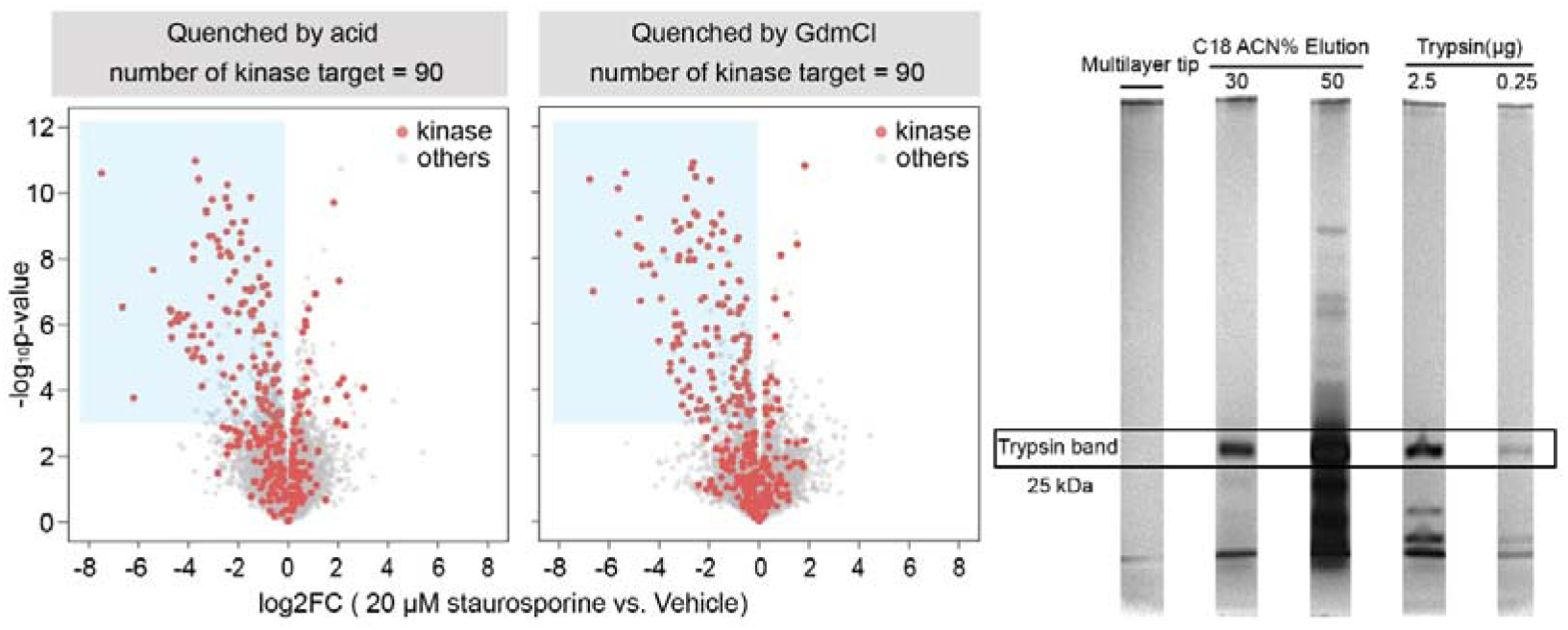
Optimization of the quenching and cleanup steps in AutoPELSA. (a) Volcano plots showing PELSA results for staurosporine obtained using acid quenching or guanidine hydrochloride quenching. The kinase targets identified under each condition are indicated above the plots. (b) Silver-stained SDS–PAGE of eluates generated from three cleanup methods: the multilayer C4–SCX–C18 tip, a 5 mg C18 tip, and a 100 mg C18 SPE column. For each method, the analyzed eluates correspond to either the 30% ACN fraction (multilayer and 5 mg C18) or the 50% ACN fraction (100 mg C18). The 25 kDa band corresponding to trypsin is indicated.

To mitigate volumetric imprecision at low pipetting volumes, we further optimized the working volumes of both trypsin and drug solutions. Collectively, these adjustments enabled full automation of the digestion process including ligand incubation, trypsinization, protein denaturation, reduction and alkylation via a liquid-handling workstation, reducing the processing time for the samples in a single 96-well plate to about 2 hours.

### Cleanup of PELSA digests using a Multilayer Tip-Column

After limited trypsinization, the PELSA digest contains small peptides, large protein fragments, residual proteins, and trypsin. For sensitive LC-MS/MS analysis of peptides, proteinaceous interferents must be removed and samples desalted. In the original PELSA workflow, these cleanup steps are conducted manually, using ultrafiltration and solid-phase extraction tips after the trypsin digestion step^10^. Together, these steps account for ∼80% of hands-on time, making them the primary source of time cost in the workflow. To address this, we consolidated protein removal and desalting into a single chromatographic step that replaces ultrafiltration and supports high-throughput operation. Recent work has evaluated single-cartridge C18 SPE as a means to concurrently deplete proteinaceous interferents and desalting samples^16^. However, evidence shows that C18 media alone are insufficient to deplete large amount of proteinaceous interferents from the mixture (Fig. 2b). And residual proteinaceous contaminants may foul the analytical column, reduce its service life, and introduce ion-suppression effects that bias MS-derived quantification.

Integration of protein removal and desalting requires effective separation of proteins from peptides, which differ in net charge and molecular size. These differences can be exploited for separation. We chose ion-exchange media due to their compatibility with tip-column implementation. However, the high ionic strength in samples (6 M guanidine HCl) outcompetes electrostatic interactions, preventing proteins and peptides from binding to SCX media (Fig. S1a). Accordingly, the ionic strength must be reduced prior to loading onto SCX. We therefore employed C4 reversed-phase media, which retained primarily through entropy-driven hydrophobic interactions^18,19^, to achieve desalting and partial separation of proteins from peptides. Building on this rationale, we designed a multilayer tip-column integrating C4, SCX, and C18 sorbents in series. C4 first retains most proteins/peptides under high ionic strength via hydrophobic interactions, enabling desalting; subsequent SCX-C18 layers further separate proteins from peptides based on charge and hydrophobicity. The separation proceeds via sequential capture, desalting, and elution across the three sorbent layers (detailed in Supplemental Materials).

We prepared stoichiometrically defined mixtures of trypsin and peptides to mimic PELSA digests and evaluated cleanup performance using two orthogonal readouts: high-sensitivity silver-stained SDS–PAGE to monitor proteinaceous contaminants in combined eluates, and absorbance at 205 nm (A205) to quantify overall peptide recovery. In contrast to a single C18 column, the multilayer column achieved superior depletion of protein fragments and residual trypsin; using the 0.25 µg trypsin band as an external reference, removal efficiency exceeded 99%, consistent with the enhanced selectivity of the multilayer design (Fig. 2b). Peptide recovery determined by A205 was ∼80.8%, comparable to recoveries obtained with conventional desalting performed without protein carryover (∼75.8%) (Fig. S1b), and thus is favorable given the multilayer sorbent architecture, which typically incurs additional loss. As a functional benchmark, staurosporine target profiling showed that the multilayer tip–based AutoPELSA identified more kinase targets than the original PELSA under identical analytical conditions (114 vs. 101; Fig. S2).

For high-throughput operation, multilayer tips were processed by positive-pressure SPE on a pneumatic manifold used in tandem with the pipetting workstation (Fig. 1b). In such systems, variability in manually packed sorbent beds can introduce backpressure heterogeneity and uneven flow that may promote tip dry-out and consequent sample loss. To mitigate these effects, a glass-fiber membrane was placed above the multilayer sorbent^20^. The membrane imposes capillary backpressure at the gas-liquid interface, helping equalize flow across tips, reducing dry-out and associated losses, and enabling reliable operation at ≤0.1 bar, sufficient to complete the multilayer tip-column procedure. The multilayer columns enable completion of a 96-well plate in ∼2 h (∼50× the throughput of the manual workflow). The tips are inexpensive (<$0.08 each), constructed from commonly available materials, and compatible with benchtop centrifugation, enhancing practicality for routine use.

In summary, AutoPELSA automates the parallel processing of 96 samples in ∼4 h, substantially increasing throughput while minimizing interference associated with single-cartridge C18 SPE desalting.

### Benchmarking AutoPELSA by identifying target proteins of staurosporine

We benchmarked AutoPELSA against the manual PELSA workflow using K562 cell lysates with eight technical replicates. AutoPELSA identified more proteins and peptides than the manual protocol (Fig. 3a). Quantitatively, the median coefficient of variation (CV) was 18.2% for AutoPELSA versus 24.0% for manual PELSA, indicating improved precision (Fig. 3b). Pearson correlation analyses also showed higher between-replicate concordance for AutoPELSA (Fig. 3c), consistent with reduced variability under automated operation.

**Figure 3.**
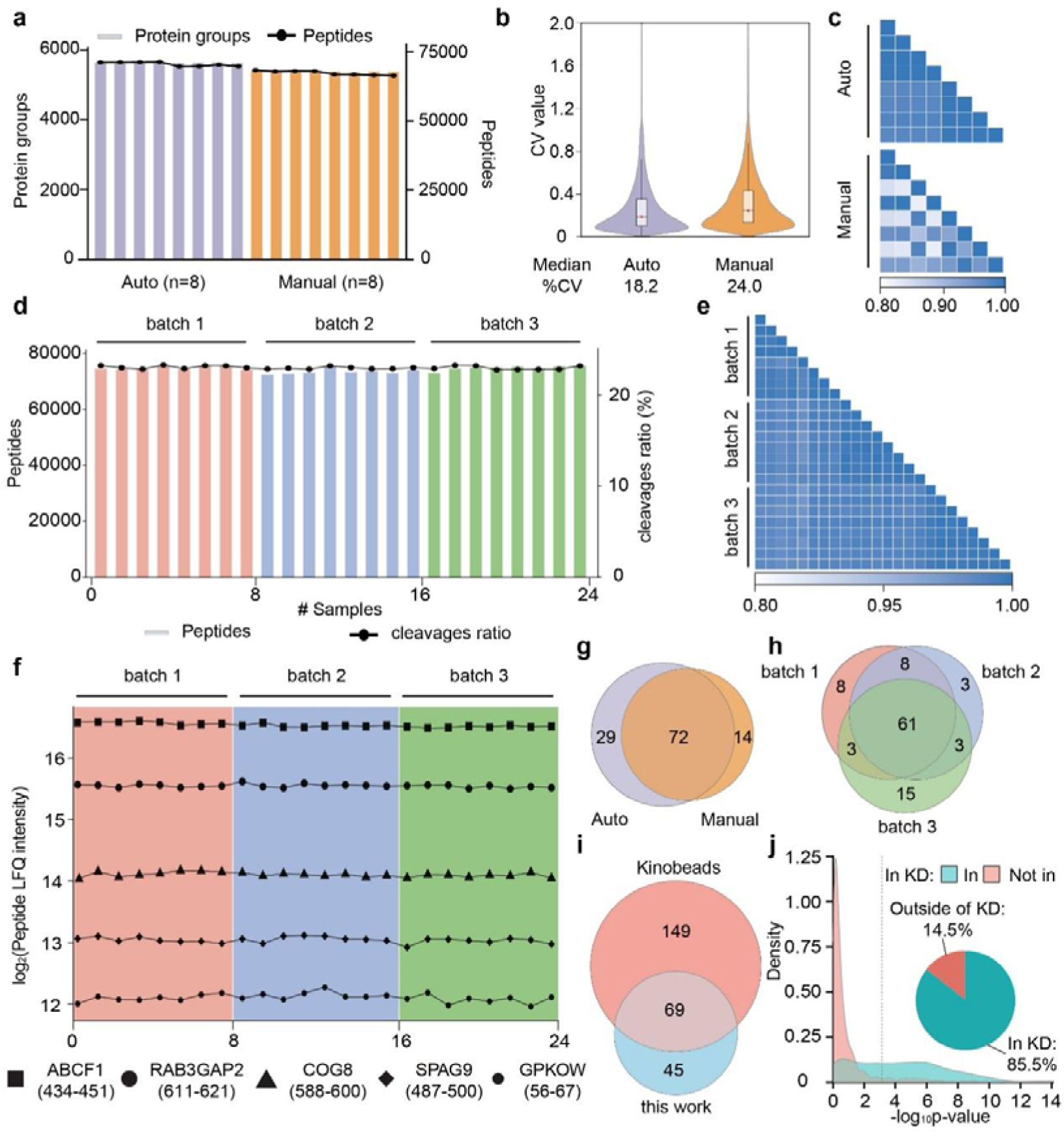
Performance assessment of the AutoPELSA workflow. (a) Numbers of identified proteins and peptides from eight technical replicates processed by AutoPELSA and manual PELSA. (b) Distributions of peptide LFQ intensity CVs for automated and manual workflows. (c) Pearson correlation matrices of peptide LFQ intensities for AutoPELSA and manual PELSA. (d) Numbers of identified peptides and percentages of peptides containing missed trypsin cleavages across three independent AutoPELSA batches. (e) Pearson correlations matrices of peptide LFQ intensities among the three batches. (f) LFQ intensity profiles of five representative peptides spanning different abundance ranges across all batches. (g) Venn diagram showing overlap of kinase targets identified by AutoPELSA and manual PELSA (each aggregated across three batches). (h) Overlap of kinase targets identified across three AutoPELSA batches. (i) Comparison of kinase targets identified by AutoPELSA and kinobeads assays. (j) Density distributions of -log_10_(p-value) for kinase-derived peptides located inside or outside annotated kinase domains. The dashed lines indicate −log_10_(p-value) cutoffs = 3. The pie charts show the locations of the kinase peptides that passed −log_10_(p-value) cutoffs.

To assess reproducibility, we processed three 96-well plates of K562 lysates (50 µg per well) on three separate days (total n = 288 samples). For each batch, eight wells were randomly selected (n = 24 total) to evaluate intra- and inter-batch performance. Across the three batches, we identified an average of 74,805 peptides, of which 32.8% contained missed trypsin cleavage sites. The missed-cleavage rate—a sensitive indicator of digestion quality in PELSA— remained stable across batches, reflects a well-controlled, reproducible trypsinization process under automated operation. The total number of identified proteins and fully cleaved peptides varied by <2% across batches, reflecting highly reproducible sample preparation and consistent identification depth (Fig. 3d). The intensity distributions of quantified peptides were highly consistent across batches (Fig. S4a). Median CVs within batches were 17.5%, 15.0%, and 13.4% for batches 1–3, respectively (Fig. S4b), demonstrating highly consistent peptide quantification within each batch. Furthermore, we selected five peptides with different LFQ intensity ranges from 2^12^ to 2^17^, and their CVs were below 4% across all three batches (Fig. 3f). For inter-batch comparisons, Pearson correlation coefficients for the 24 profiled samples were >0.94 (Fig. 3e) with intra-batch correlations >0.95, consistent with high quantitative reproducibility. Collectively, these results demonstrate that AutoPELSA provides high intra- and inter-batch reproducibility in sample preparation.

To further evaluate target-identification performance, we profiled staurosporine across three independent AutoPELSA batches performed on different days. Relative to manual PELSA, AutoPELSA identified 15 additional kinase targets, while 72 kinase targets were shared by both methods (Fig. 3g). Across the three automated batches, 80, 75, and 82 kinase targets were identified, with 61 kinase targets consistently detected in all runs (Fig. 3h), indicating stable target recovery across batches.

To assess reliability, we compared AutoPELSA results with kinobeads pull-down assays, a widely used method for kinase target identification. A total of 69 kinase targets were commonly detected by both approaches, whereas kinobeads exclusively identified 149 targets and AutoPELSA exclusively identified 45(Fig. 3i). This difference may because: kinobeads data were compiled across multiple cell lines and experimental replicates, while the AutoPELSA results reported here reflect a single-cell-line measurement. Among the identified 114 kinase targets, 85.5% of the peptides with -log_10_(p-value) > 3 were located within kinase domains (Fig. 3j), further supporting the accuracy of target assignment by PELSA.

### AutoPELSA allows efficient identification of low affinity ligand targets

To evaluate the capability of AutoPELSA to detect targets of low-affinity ligands, we applied our method to α-ketoglutarate (α-KG), a metabolite known to bind proteins with low affinity^10, 21^. Guided by previous reports showing that 5 mM α-KG supports stem cell viability^22^, we selected two concentration points-2 mM and 10 mM-for target identification in K562 cells. AutoPELSA identified 41 significantly ligand-stabilized proteins at 2 mM α-KG (-log_10_(p-value) > 3.4, logLFC < −0.5; Bayesian t-test), 30 of which correspond to known targets. At 10 mM, 424 ligand-stabilized proteins were detected, including 29 previously reported targets (Fig. 4a). Of the 32 proteins significantly stabilized at both α-KG concentrations, 26 corresponded to previously reported targets. In contrast, 392 and 9 proteins were uniquely stabilized at 10 mM and 2 mM, respectively (Fig. 4b). The higher proportion of known targets detected at 2 mM α-KG may reflect the closer alignment of this concentration with physiological conditions, facilitating capture of endogenous interaction partners. This demonstrates that AutoPELSA effectively resolves ligand-induced local stability changes, even for low-affinity interactions. Among the significantly stabilized peptides, 21 proteins containing annotated α-KG–binding domains were identified, and 16 exhibited >20% coverage of these domains (Fig. 4d), indicating that AutoPELSA can capture stability shifts that localize to ligand-binding regions. KEGG pathway enrichment analysis revealed significant accumulation of the identified αKG-binding proteins in metabolic pathways, including IDH1 and IDH2 from the tricarboxylic acid cycle and ACLY, which bridges carbohydrate and lipid metabolism (Fig. 4c). Beyond core metabolism, AutoPELSA also detected members of the Jumonji C domain–containing (JMJD) protein family, such as JMJD4 and JMJD6, which are well known as epigenetic regulators that act predominantly as histone lysine or arginine demethylases^23, 24^. Notably, the local stability profiles, generated by PELSA Decipher^25^, precisely revealed α-KG interactions to the conserved JmjC catalytic domain (Fig. 4e), demonstrating domain-level engagement consistent with its proposed role in epigenetic regulation.

**Figure 4.**
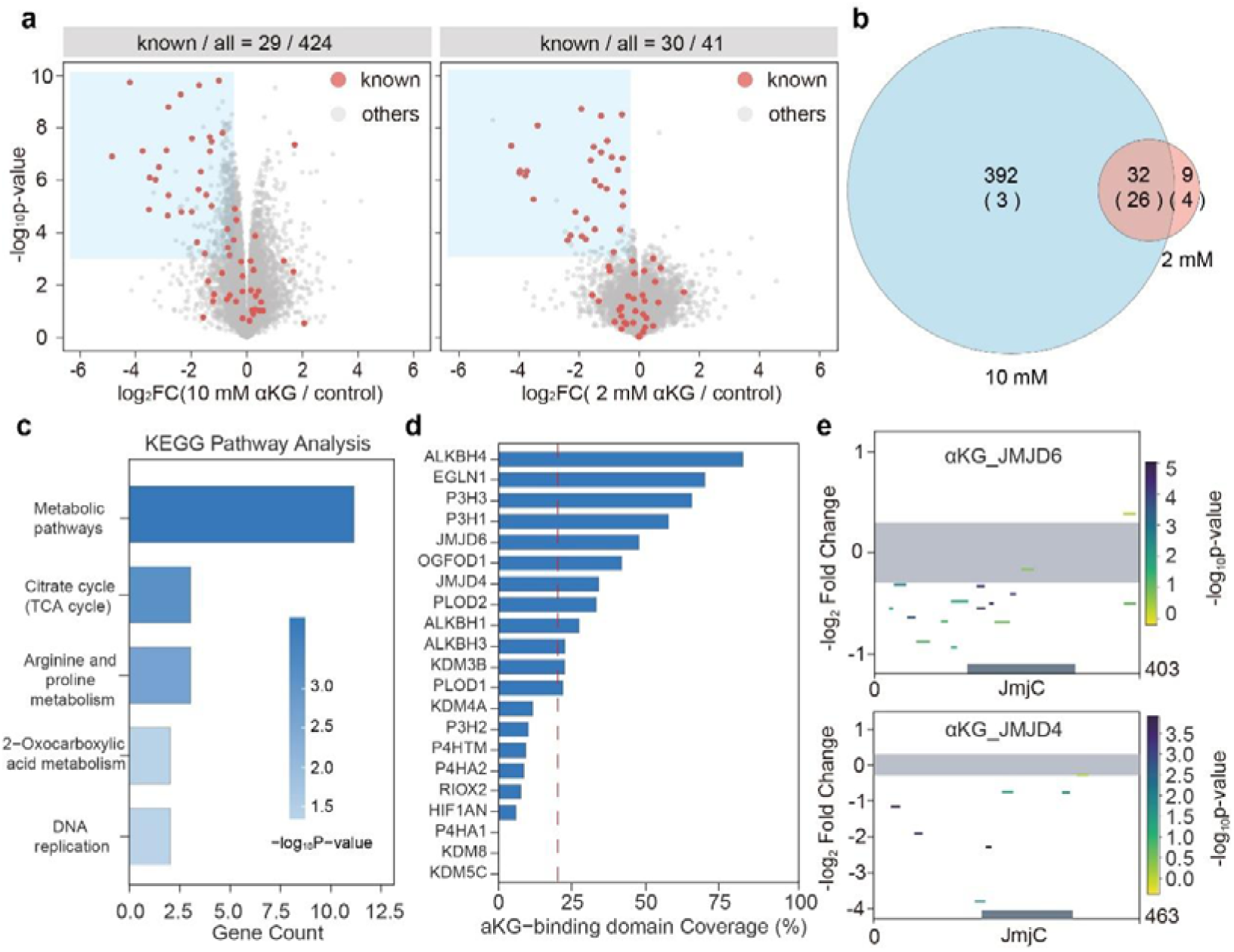
Identification of α-ketoglutarate (αKG) targets using AutoPELSA. (a) Volcano plots of proteins showing αKG-induced stabilization at 10 mM and 2 mM (four replicates each). Previously reported αKG-binding proteins are highlighted. Statistical thresholds used for significance assessment are shown in the shaded region. (b) Overlap of stabilized proteins identified at the two αKG concentrations. Numbers in parentheses indicate previously known αKG targets. (c) KEGG pathway enrichment analysis of proteins stabilized at 2 mM αKG. (d) Peptide coverage within annotated αKG-binding domains for 21 known targets (-log_10_(p-value) > 3). The vertical dashed line marks 20% domain coverage. (e) Local stability profiles of JMJD6 and JMJD4 peptides mapped across the conserved JmjC domain under αKG-treated conditions. The local stability profiles were generated by PELSA Decipher

### AutoPELSA Enables Measurement of Ligand-Protein Interaction Affinities in Cell Lysates

Traditional ligand–protein interaction assays often require recombinant protein expression and domain truncation, limiting their physiological relevance^26–28^. In contrast, PELSA preserves the native state of proteins in cell lysates and is therefore well-suited for affinity profiling. However, such studies require dose-dependent measurements across multiple conditions, making automation essential. We therefore employed AutoPELSA to perform dose-dependent interaction analyses with high throughput and reproducibility.

To exploit the ability of AutoPELSA to measure ligand–protein interactions across thousands of proteins in parallel, we designed a mixed-drug assay to simultaneously profile multiple ligands in a single experiment. By staggering the concentration series of each ligand, distinct dose–response signatures can be inferred directly from the mixed solutions, allowing EC values to be estimated for each ligand–target pair and enabling discrimination of target classes based on their binding affinities. Four ligands with distinct target classes were selected: methotrexate (DHFR), lapatinib (EGFR), ganetespib (HSP90 family), and rapamycin (FKBP family). Serial dilutions of each ligand were staggered and combined to generate mixed-drug solutions (Fig. 5a), enabling parallel dose–response profiling for all four ligands within a single proteomic run.

**Figure 5.**
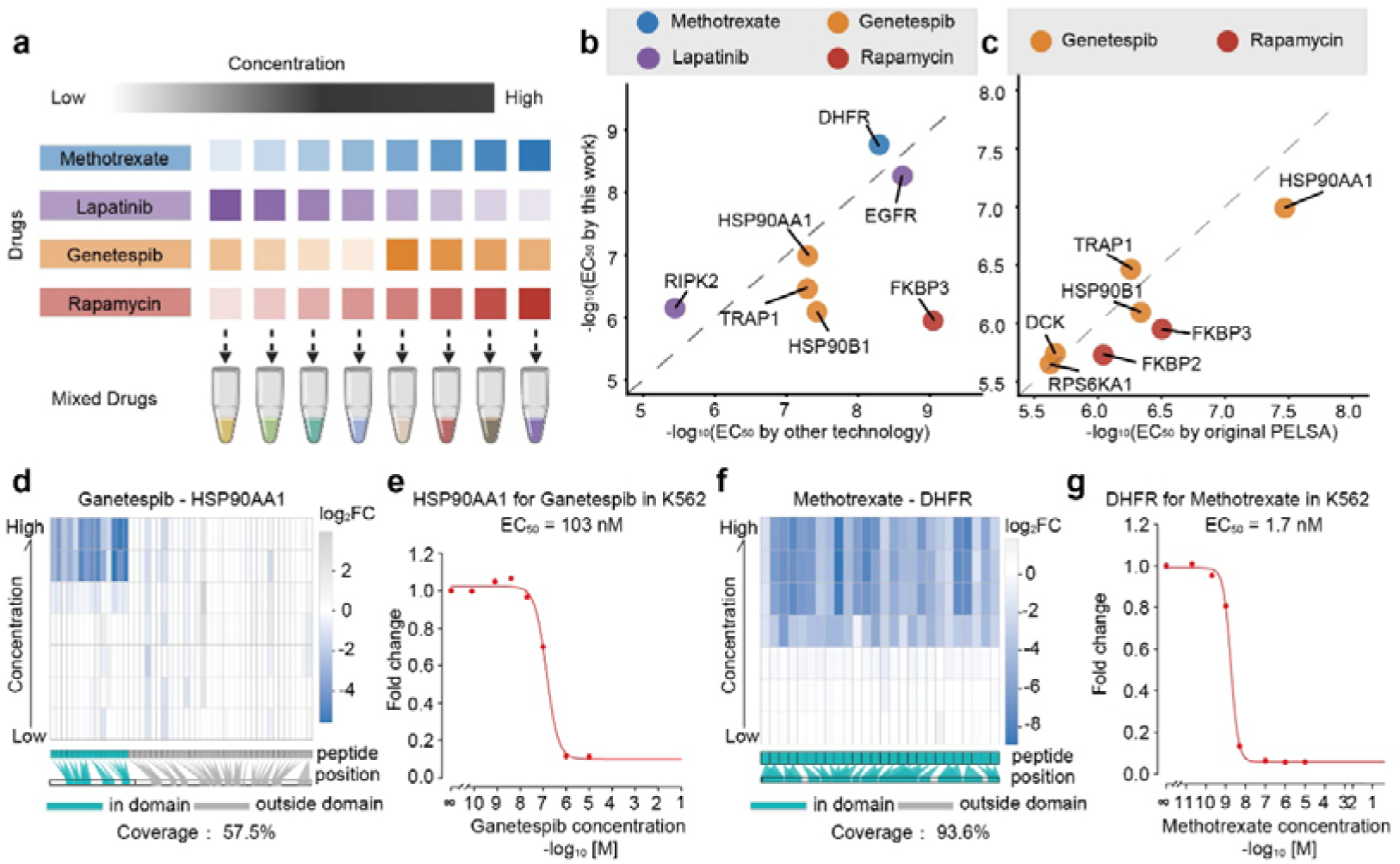
Multiplexed affinity profiling using mixed-ligand dose–response AutoPELSA. (a) Experimental design for staggered concentration series of four ligands (rapamycin, ganetespib, methotrexate, lapatinib) allowing simultaneous dose–response measurement within a single proteomic run. (b) Comparison of ECOO values obtained from AutoPELSA with measurements by other techniques. (c) Comparison of ECOO values obtained from AutoPELSA with previously reported values from the original PELSA workflow. (d,f) Heatmaps showing region-specific stabilization of HSP90AA1 (ganetespib) and DHFR (methotrexate). Peptides mapped to annotated ligand-binding regions are shown in cyan, and peptides outside these regions are shown in gray. (e,g) Representative dose–response curves for ganetespib–HSP90AA1 and methotrexate–DHFR interactions derived from AutoPELSA. ECOO values for dose dependent PELSA experiments were computed by PELSA Decipher.

The mixed-drug dose–response strategy assigned each ligand a staggered concentration sequence (Fig. 5a), enabling deconvolution of binding affinities from combined samples. Across the quantified ligand–target pairs, ECLL values derived from AutoPELSA showed strong agreement with those obtained using orthogonal affinity measurement approaches, including enzyme activity inhibition assays^29^, fluorescence polarization^30^, ATP-site–dependent competition binding assays^31^, and rapamycin affinity matrix–based methods^32^ (Fig. 5b). Most targets clustered near the identity line, with the majority of deviations remaining within one order of magnitude. Above data are consistent with published PELSA data^10^ (Fig. 5c), indicating that this implementation retains the fundamental principles of PELSA while enabling multiplexed analysis. Notably, targets with higher domain coverage (e.g., DHFR and EGFR) tended to align more closely with known affinities, suggesting that domain-level resolution enhances the reliability of affinity estimation. Furthermore, analyses of representative targets demonstrated region-specific stabilization responses. As shown in Fig. 5d-g, ligand-induced stabilization was spatially localized and consistently observed only within annotated binding domains, while non-binding regions remained largely unchanged, indicating that local stability shifts detected by AutoPELSA are highly selective, supporting the mechanistic specificity of the interaction profiles. Collectively, these results demonstrate that AutoPELSA can recover binding-region–specific stability changes and determine ligand affinities for multiple targets in a single proteomic experiment, while retaining the ability to resolve interactions at domain-level resolution under native conditions.

## Conclusion

In this study, we developed AutoPELSA, an automated workflow for large-scale identification of ligand–protein interactions under native conditions. The workflow integrates two functionally coordinated modules: a fully automated limited trypsinization process executed on a liquid-handling workstation, and a custom C4-SCX-C18 multilayer tip-column that enables simultaneous removal of partially digested protein fragments, depletion of residual trypsin, and peptide desalting. This design consolidates multiple manual procedures into a single chromatographic step, allowing the entire workflow to be completed within four hours, thereby achieving a >20-fold increase in throughput compared to the manual protocol. Automated processing yielded high reproducibility and quantitative consistency, enabling robust recovery of ligand–protein interactions. Notably, AutoPELSA successfully identified targets of low-affinity ligands while localizing the binding regions, revealing localized stability changes within interaction regions. Furthermore, through a mixed-ligand dose–response strategy, AutoPELSA simultaneously quantified binding affinities for multiple ligands in a single experiment, demonstrating its suitability for multiplexed affinity profiling. Overall, AutoPELSA represents a high-throughput, modification-free platform for proteome-wide interrogation of ligand–protein interactions in their native biochemical context. By enabling peptide-level binding affinity measurements at scale, it provides a practical route to deciphering the mechanisms of action of drug candidates, supporting systematic mapping of ligand–target landscapes that can guide rational drug development.

## Acknowledgments

We acknowledge the financial supports from the National Key Research and Development Program of China (2021YFA1302600), the National Natural Science Foundation of China (22437007), Dalian Science and Technology Innovation Fund (2023JJ11CG006), the Innovation Program of Science and Research from the DICP, CAS (DICP I202109).

## Data Availability

The main data supporting the results in the study are available within the paper and its supplementary information. The raw data has been uploaded onto JPOST Repository^33^. The accession number PXD072751 for ProteomeXchange and JPST004294 for jPOST (URL: https://repository.jpostdb.org/preview/1981498290695e3e980e28b, Access key: 4438)

## TOC

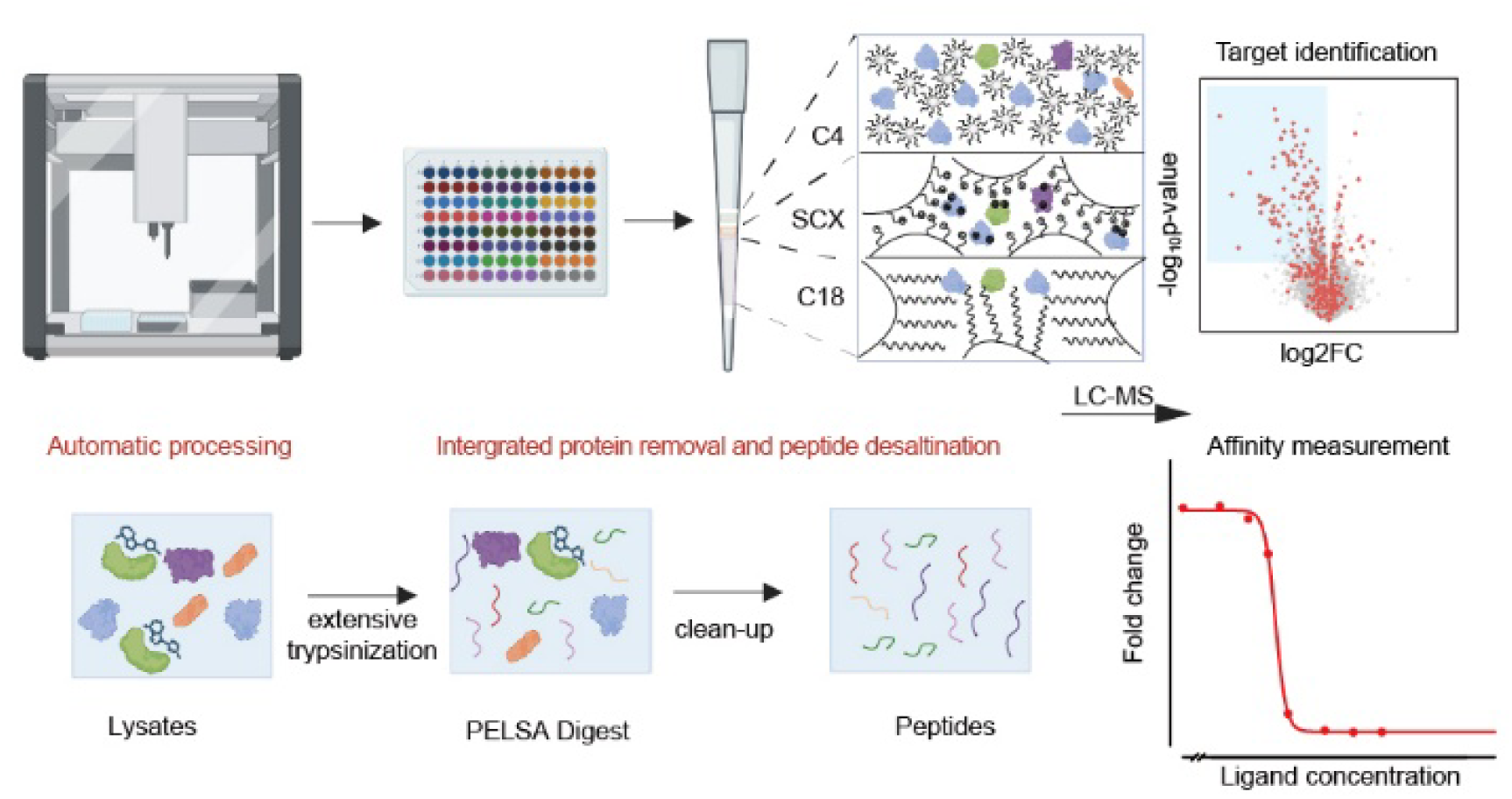

## Materials and Methods

### Chemicals and Reagents

Lapatinib, Methotrexate, Rapamycin, Ganetespib, α-ketoglutarate, and Staurosporine were purchased from Selleck. Other chemicals were from Sigma-Aldrich unless otherwise specified.

### Cell Culture

K562 (ATCC CCL-243) cells were cultured in IMDM with 10% BS, 1% penicillin, and 1% streptomycin at 37 °C and 5% COL. Cells were harvested by centrifugation, washed with PBS, and stored at −80 °C.

### Multi-layer Tip Fabrication

Tips were sequentially packed with C18 (SepPak, Waters), SCX (MC30-SP, Sepax), and C4 resins (XB-C4, Welch), separated by hydrophobic filters (BF014-12-20, biocomma) or glass membranes (Grade GF/C Glass Microfibre Filter, Cytiva). Detailed specifications are provided in Supplemental Materials.

### Manual PELSA

Manual PELSA was performed following our previously published protocols^10^.

### Trypsin Removal Assay

The efficiency of SCX, C18, and multi-layer tips in removing trypsin was assessed using BCA assay or absorbance at 205 nm. Detailed protocols are described in Supplemental Materials.

### AutoPELSA Workflow

Automated PELSA was performed using an Opentrons FLEX Robot with homemade multi-layer tips. The workflow included: (1) Ligand incubation (30 min, room temperature, 4 replicates). (2) Native trypsinization (40 s, 37 °C). (3) Protein denaturation and peptide reduction/alkylation. (4) Peptide acidification and cleanup using multi-layer tips. All ligand concentrations and cell lines are listed in the Supplemental Table. Detailed robotic deck layouts, volumes, and solutions are described in Supplemental Materials.

### LC-MS/MS Analysis

Peptides were analyzed on an Orbitrap Exploris 480 with micro-flow LC. Separation was performed on a C18 column with linear gradients; DIA acquisition was used. Full and MS/MS scan parameters are detailed in Supplemental Materials.

### Peptide Identification and Quantification

Data were processed using Spectronaut directDIA (Biognosys AG, v.19) against the UniProt human database (2024 release). Detailed search parameters provided in the Supporting Information.

### Statistical Analysis

Peptide intensities were analyzed in R using the limma package. Two-sided empirical Bayes moderated t-tests were performed to compare ligand-treated versus control samples. Log_2_FC represents stability changes; -log_10_(p-value) indicates significance. Figure 4e shows the local stability profile of the bound domains, generated using PELSA Decipher^25^. EC_50_ values were determined by sigmoidal curve fitting in PELSA Decipher.

## References

1. Klingenberg, M., Ligand-protein interaction in biomembrane carriers. The induced transition fit of transport catalysis. Biochemistry 2005, 44 (24), 8563–8570.

2. Gottschalk, K. E.; Kessler, H., The structures of Integrins and integrin-ligand complexes: Implications for drug design and signal transduction. Angewandte Chemie-International Edition 2002, 41 (20), 3767–3774.

3. Huang, Z. W., Structural chemistry and therapeutic intervention of protein-protein interactions in immune response, human immunodeficiency virus entry, and apoptosis. Pharmacol. Ther. 2000, 86 (3), 201–215.

4. Kim, P.; Zhao, J. F.; Lu, P. Y.; Zhao, Z. M., mutLBSgeneDB: mutated ligand binding site gene DataBase. Nucleic Acids Res. 2017, 45 (D1), D256–D263.

5. Wang, H. P.; Xiong, R. Y.; Lai, L. H., Rational drug design targeting intrinsically disordered proteins. Wiley Interdisciplinary Reviews-Computational Molecular Science 2023, 13 (6).

6. Lazar, T.; Connor, A.; DeLisle, C. F.; Burger, V.; Tompa, P., Targeting protein disorder: the next hurdle in drug discovery. Nature Reviews Drug Discovery 2025.

7. Tarantino, P.; Ricciuti, B.; Pradhan, S. M.; Tolaney, S. M., Optimizing the safety of antibody-drug conjugates for patients with solid tumours. Nature Reviews Clinical Oncology 2023, 20 (8), 558–576.

8. Meissner, F.; Geddes-McAlister, J.; Mann, M.; Bantscheff, M., The emerging role of mass spectrometry-based proteomics in drug discovery. Nature Reviews Drug Discovery 2022, 21 (9), 637–654.

9. Feng, Y.; De Franceschi, G.; Kahraman, A.; Soste, M.; Melnik, A.; Boersema, P. J.; de Laureto, P. P.; Nikolaev, Y.; Oliveira, A. P.; Picotti, P., Global analysis of protein structural changes in complex proteomes. Nat Biotechnol 2014, 32 (10), 1036–44.

10. Li, K. J.; Chen, S. J.; Wang, K. Y.; Wang, Y.; Xue, L. J.; Ye, Y. Y.; Fang, Z.; Lyu, J. W.; Zhu, H. Y.; Li, Y. N.; Yu, T.; Yang, F.; Zhang, X. L.; Guo, S. Q.; Ruan, C. F.; Zhou, J. H.; Wang, Q.; Dong, M. M.; Luo, C.; Ye, M. L., A peptide-centric local stability assay enables proteome-scale identification of the protein targets and binding regions of diverse ligands. Nature Methods 2025, 22 (2).

11. Wang, K. Y.; Li, Y. A.; Ma, Y. N.; Ye, M. L., Probing ligand-induced local stability shifts: A sensitive approach to identify target proteins and binding sites at the proteomic scale. Curr. Opin. Chem. Biol. 2025, 87.

12. Wang, K.; Li, Y.; Ma, Y.; Ye, M., Probing ligand-induced local stability shifts: A sensitive approach to identify target proteins and binding sites at the proteomic scale. Curr. Opin. Chem. Biol. 2025, 87.

13. Wu, Q.; Zheng, J.; Sui, X.; Fu, C.; Cui, X.; Liao, B.; Ji, H.; Luo, Y.; He, A.; Lu, X.; Xue, X.; Tan, C. S. H.; Tian, R., High-throughput drug target discovery using a fully automated proteomics sample preparation platform. Chemical Science 2024, 15 (8), 2833–2847.

14. Schär, S.; Räss, L.; Malinovska, L.; Savickas, S.; Cavallo, F.; Below, C.; Tognetti, M.; Shichkova, P.; Gourdet, B.; Robles, G.; Iu, L.; Vowinckel, J.; Feng, Y.; Hjerpe, R.; Bruderer, R.; Reiter, L., A Flexible End-to-End Automated Sample Preparation Workflow Enables Standardized and Scalable Bottom-up Proteomics. Analytical Chemistry 2025.

15. Arad, M.; Frey, C.; Balagtas, R.; Hare, R.; Ku, K.; Jereb, D.; Nestman, Z.; Sidhu, A.; Shi, Y.; Fordwour, O.; Moon, K.-M.; Foster, L. J.; Ghafourifar, G., Development of an Automated, Ultra-Rapid Bottom-Up Proteomics Workflow Utilizing Alginate-Based Hydrogels. Analytical Chemistry 2024.

16. Li, K.; Potel, C. M.; Becher, I.; Hüttmann, N.; Garrido-Rodriguez, M.; Schwarz, J.; Savitski, M. M., High-throughput peptide-centric local stability assay extends protein-ligand identification to membrane proteins, tissues, and bacteria. bioRxiv 2025, 2025.04.28.650974.

17. Malthouse, J. P. G., Kinetic Studies of the Effect of pH on the Trypsin-Catalyzed Hydrolysis of *N*-α-benzyloxycarbonyl-L-lysine-*p*-nitroanilide: Mechanism of Trypsin Catalysis. Acs Omega 2020, 5 (10), 4915–4923.

18. 18. Hydrophobic Interaction Chromatography of Proteins. Lcgc Europe 2019, 32 (3), 160–160.

19. Queiroz, J. A.; Tomaz, C. T.; Cabral, J. M. S., Hydrophobic interaction chromatography of proteins. Journal of Biotechnology 2001, 87 (2), 143–159.

20. Zougman, A.; Selby, P. J.; Banks, R. E., Suspension trapping (STrap) sample preparation method for bottom- up proteomics analysis. Proteomics 2014, 14 (9), 1006–1010.

21. Piazza, I.; Kochanowski, K.; Cappelletti, V.; Fuhrer, T.; Noor, E.; Sauer, U.; Picotti, P., A Map of Protein-Metabolite Interactions Reveals Principles of Chemical Communication. Cell 2018, 172 (1-2), 358-+.

22. Cui, Z.; Li, J.; Li, C.; Deng, S.; Liu, W.; Lei, T.; Cao, J.; Wang, Z.; Wang, X.; Ma, S.; Zhu, Y.; Yang, H.; Chen, P., Identifying the target, mechanism, and agonist of α-ketoglutaric acid in delaying mesenchymal stem cell senescence. Cell Reports 2025, 44 (7).

23. Wang, P.; Chen, L.-L.; Xiong, Y.; Ye, D., Metabolite regulation of epigenetics in cancer. Cell Reports 2024, 43 (10).

24. Wang, M.; Xue, J.; Hong, W.; Chen, S.; Shi, H., JMJD family proteins in cancer and inflammation. Signal Transduction and Targeted Therapy 2022, 7 (1).

25. Zhu, H.; Wang, K.; Li, K.; Fang, Z.; Zhou, J.; Xue, L.; Ye, M., PELSA-Decipher: A Software Tool for the Processing and Interpretation of Ligand-Protein Interaction Data Sets Acquired by PELSA. Journal of Proteome Research 2025, 24 (9), 4623–4630.

26. Du, X.; Li, Y.; Xia, Y.-L.; Ai, S.-M.; Liang, J.; Sang, P.; Ji, X.-L.; Liu, S.-Q., Insights into Protein–Ligand Interactions: Mechanisms, Models, and Methods. International Journal of Molecular Sciences 2016, 17 (2), 144.

27. Serdiuk, T.; Fleischmann, Y.; Ghosh, D.; Delparente, A.; Reber, V.; Frey, L.; Rhyner, D.; Gerez, J.; Volkmar, N.; Kralickova, L.; Mas, G.; Doerig, C.; Hiller, S.; Schibli, R.; Mu, L.; Picotti, P.; Riek, R., Alpha-synuclein interacts with regulators of ATP homeostasis in mitochondria. Nature Communications 2025, 16 (1).

28. Patricelli, M. P.; Nomanbhoy, T. K.; Wu, J. Y.; Brown, H.; Zhou, D.; Zhang, J. M.; Jagannathan, S.; Aban, A.; Okerberg, E.; Herring, C.; Nordin, B.; Weissig, H.; Yang, Q. K.; Lee, J. D.; Gray, N. S.; Kozarich, J. W., In Situ Kinase Profiling Reveals Functionally Relevant Properties of Native Kinases. Chem. Biol. 2011, 18 (6), 699–710.

29. Tsukamoto, T.; Haile, W. H.; McGuire, J. J.; Coward, J. K., Synthesis and Biological Evaluation of Nα-(4-Amino-4-deoxy-10-methylpteroyl)-dl-4,4-difluoroornithine. J. Med. Chem. 1996, 39 (13), 2536–2540.

30. Taldone, T.; Patel, P. D.; Patel, M.; Patel, H. J.; Evans, C. E.; Rodina, A.; Ochiana, S.; Shah, S. K.; Uddin, M.; Gewirth, D.; Chiosis, G., Experimental and Structural Testing Module to Analyze Paralogue-Specificity and Affinity in the Hsp90 Inhibitors Series. J. Med. Chem. 2013, 56 (17), 6803–6818.

31. Karaman, M. W.; Herrgard, S.; Treiber, D. K.; Gallant, P.; Atteridge, C. E.; Campbell, B. T.; Chan, K. W.; Ciceri, P.; Davis, M. I.; Edeen, P. T.; Faraoni, R.; Floyd, M.; Hunt, J. P.; Lockhart, D. J.; Milanov, Z. V.; Morrison, M. J.; Pallares, G.; Patel, H. K.; Pritchard, S.; Wodicka, L. M.; Zarrinkar, P. P., A quantitative analysis of kinase inhibitor selectivity. Nature Biotechnology 2008, 26 (1), 127–132.

32. Fretz, H.; Albers, M. W.; Galat, A.; Standaert, R. F.; Lane, W. S.; Burakoff, S. J.; Bierer, B. E.; Schreiber, S. L., RAPAMYCIN AND FK506 BINDING-PROTEINS (IMMUNOPHILINS). Journal of the American Chemical Society 1991, 113 (4), 1409–1411.

33. Okuda, S.; Watanabe, Y.; Moriya, Y.; Kawano, S.; Yamamoto, T.; Matsumoto, M.; Takami, T.; Kobayashi, D.; Araki, N.; Yoshizawa, A. C. J. N. a. r., jPOSTrepo: an international standard data repository for proteomes. 2017, 45 (D1), D1107–D1111.

