## supplementary1 for "AutoPELSA: an automated sample preparation system for proteome-wide identification of target proteins of diverse ligands"

Corresponding authors:

**Supplementary methods:**

**Fabrication of the multi-layer tip**

Multi-layer tips were constructed by sequentially packing 5 mg of C18 resin (SepPak, Waters), 2.5 mg of SCX resin (MC30-SP, Sepax), and 2.5 mg of C4 resin (XB-C4, Welch) into a 200 μL pipette tip (T-200-Y, Axygen) from bottom to top. A hydrophobic filter (BF014-12-20, biocomma) was placed at the tip bottom, and three glass fiber plugs (Grade GF/C Glass Microfibre Filter, Cytiva) were inserted at the top to separate and stabilize the resin layers. Hydrophobic filters or glass membranes were optionally used to further separate distinct filler layers.

**Assay the effciency of trypsin removing with different types of tips**
SCX tips were prepared by packing 5 mg of SCX resin. After activation with 30% methanol and equilibration with potassium citrate buffer (PCB, pH 3.0), samples containing 100 µg of trypsin in 10 mM PCB (pH 3.0) were loaded. The tips were subsequently washed with PCB. Bound proteins were eluted with 1 M NaCl in PCB and the eluates were quantified by BCA assay.

Multi-layer tips were prepared as described above. For sample processing, a complex mixture containing 50 µg of trypsin and 20 µg of PELSA-derived peptides in 8 M guanidine hydrochloride (pH 3.0) was loaded. The tip was activated with methanol, equilibrated with 0.1% trifluoroacetic acid (TFA), and washed after sample loading. Peptides were eluted with 30% acetonitrile (ACN)/0.1% TFA, the tip was then re-equilibrated with TFA, treated with 1 M NaCl in potassium citrate buffer (PCB), and washed again. Proteins retained after the peptide elution were subsequently eluted with a second step of 30% ACN/0.1% TFA. The two eluates were combined, and the peptide content in the combined solution was quantified by measuring the absorbance at 205 nm using a NanoDrop spectrophotometer (Thermo Fisher Scientific). The protein content in the combined eluate was analyzed by SDS–PAGE followed by silver staining(P0017S, Beyotime).

**Ligand concentrations and cell lines for AutoPELSA**

The following ligand concentrations and cell lines are used: Staurosporine (20 μM, K562), α-KG (2 mM 10mM, K562), mixed drug1 (0 nM methotrexate, 10 μM lapatinib, 4 nM ganetespib, 20 nM rapamycin, K562), mixed drug2 (0.02 nM methotrexate, 1 μM lapatinib, 0.8 nM ganetespib, 100 nM rapamycin, K562), mixed drug3 (0.2 nM methotrexate, 0.1 μM lapatinib, 0.08 nM ganetespib, 1 μM rapamycin, K562), mixed drug4 (1 nM methotrexate, 20 nM lapatinib, 0 nM ganetespib, 10 μM rapamycin, K562), mixed drug5 (5 nM methotrexate, 4 nM lapatinib, 10 μM ganetespib, 0 nM rapamycin, K562), mixed drug6 (0.1 μM methotrexate, 0.8 nM lapatinib, 1 μM ganetespib, 0.08 nM rapamycin, K562), mixed drug7 (1 μM methotrexate, 0.08 nM lapatinib, 0.1 μM ganetespib, 0.8 nM rapamycin, K562), mixed drug8 (10 μM methotrexate, 0 nM lapatinib, 20 nM ganetespib, 4 nM rapamycin, K562).

**LC-MS/MS analysis**

LC-MS/MS settings for data-independent acquisition (DIA) analysis. The samples were analyzed on Orbitrap Exploris 480 coupled with a micro-flow LC system (Ultimate 3000 RSLC microsystem, Dionex). 10 μg peptides were separated on a commercial 15 cm x 1 mm i.d. column (ACQUITY UPLC Peptide CSH C18 Column, 130 Å, 1.7 µm; Waters). Binary buffers (A, 0.1% FA; B, 80% ACN and 0.1% FA) were used. For samples displayed figure.3a-h, peptides were separated by linear gradients from 6% B to 32% B for 48 min followed by a linear increase to 45% B in 5.5 min at the flow of 50 μL/min. For other samples, peptides were separated by linear gradients from 6% B to 32% B for 80 min followed by a linear increase to 45% B in 14 min at the flow of 50 μL/min. Full MS scans were acquired at 120,000 resolution (m/z = 200) spanning from m/z 350 to 1400 with the automatic gain control (AGC) target set to 3e6 and a maximum injection time (IT) of 45 ms. MS/MS scans were acquired in a DIA mode with a resolution of 30,000 (m/z = 200). A total of 24 DIA segments were acquired ranging from m/z 400 to 1,000 at a resolution of 30,000 (AGC target of 2e6 and auto maximum IT) with the normalized collision energy (NCE) of 30%. The first mass was fixed at m/z of 300.

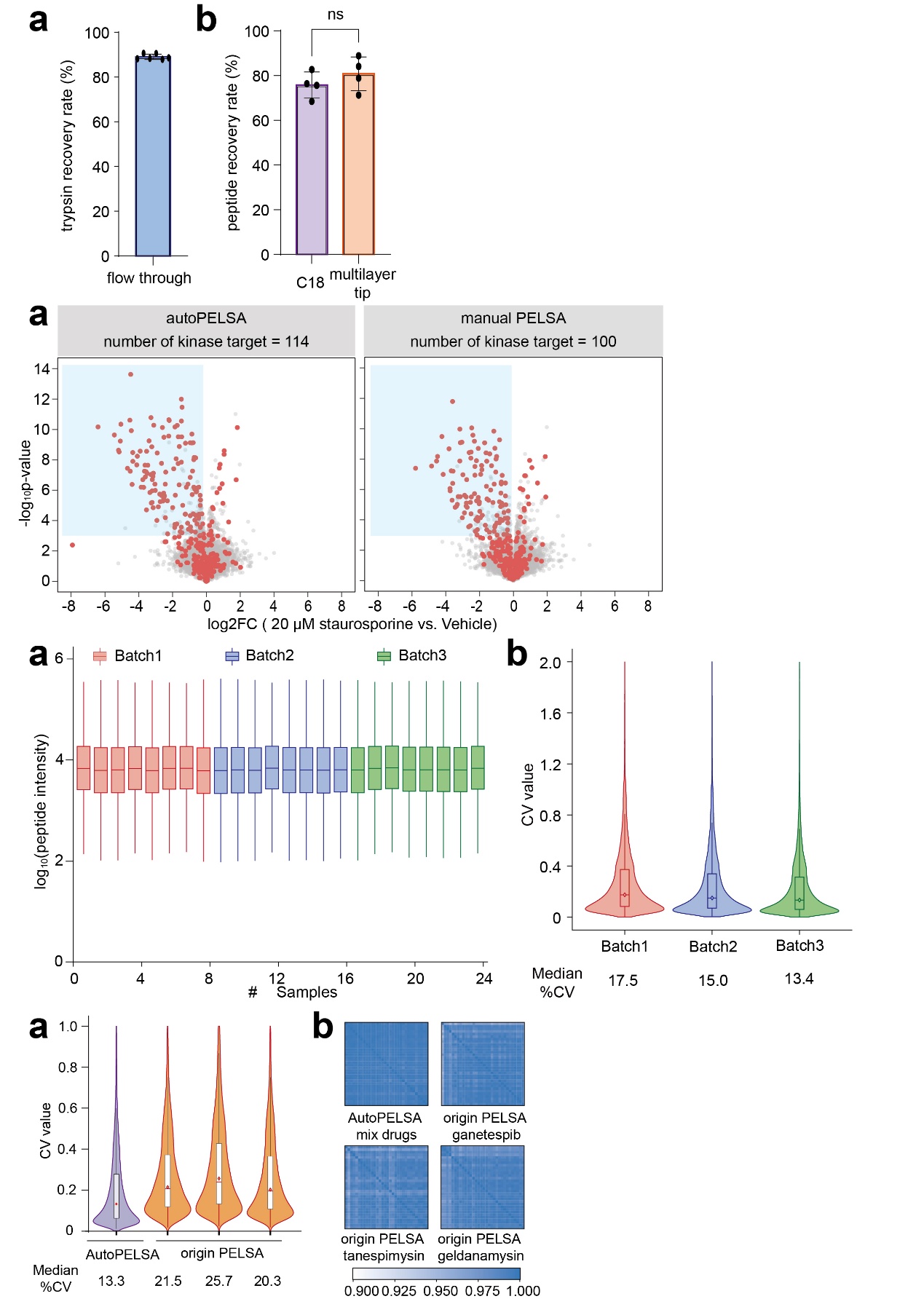

Figure S1. Separation performance of different types of tips. (a) Trypsin percentage in flow-through (SCX tip, BCA assay) (b) PELSA peptides recovery rate in eluate (C18 tip and multilayer tip, absorbance at 205 nm)

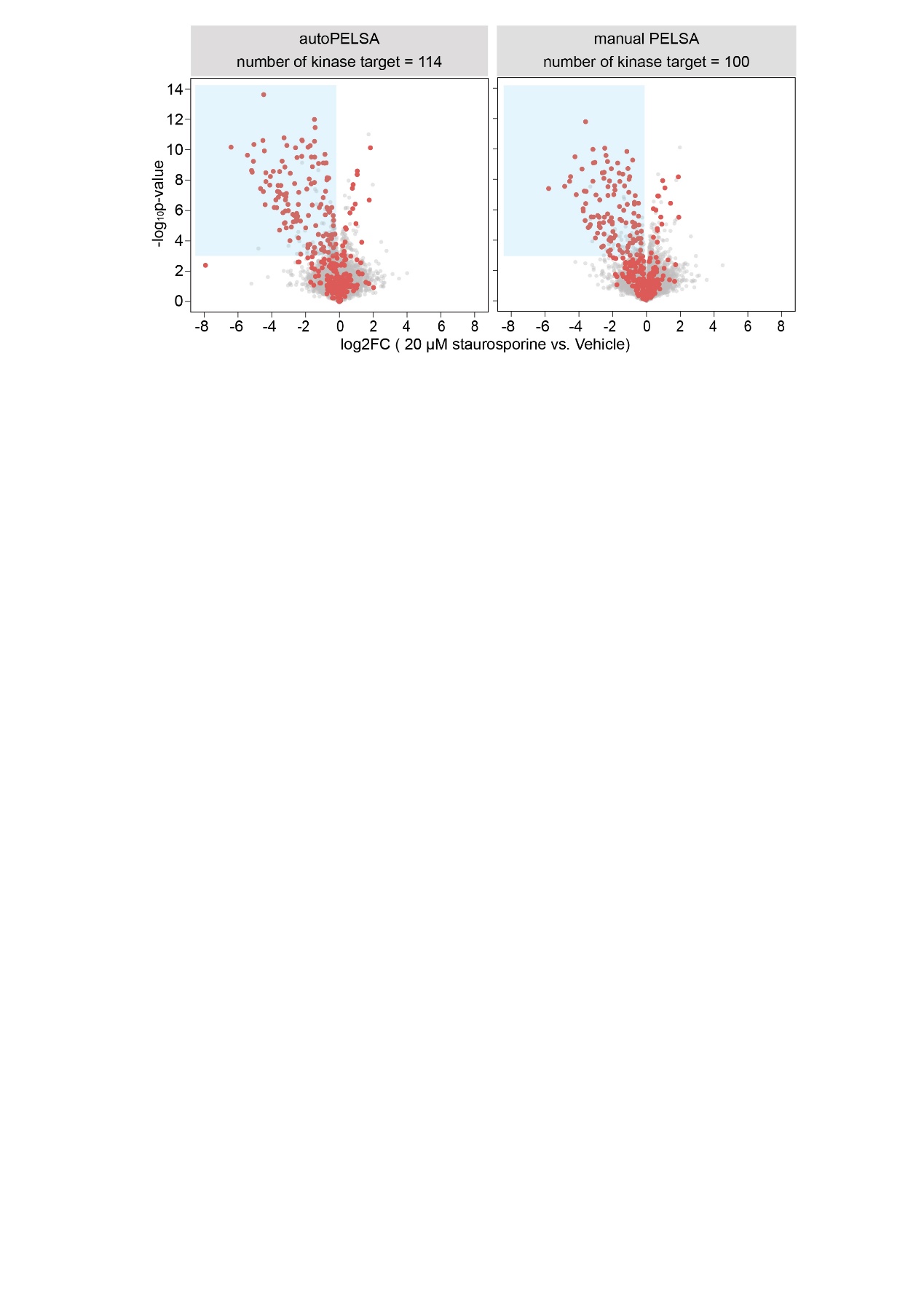

Figure S2. Comparison of identification performance between AutoPELSA and manual PELSA. Volcano plot of identified kinase targets of staurosporine using AutoPELSA and manual PELSA.

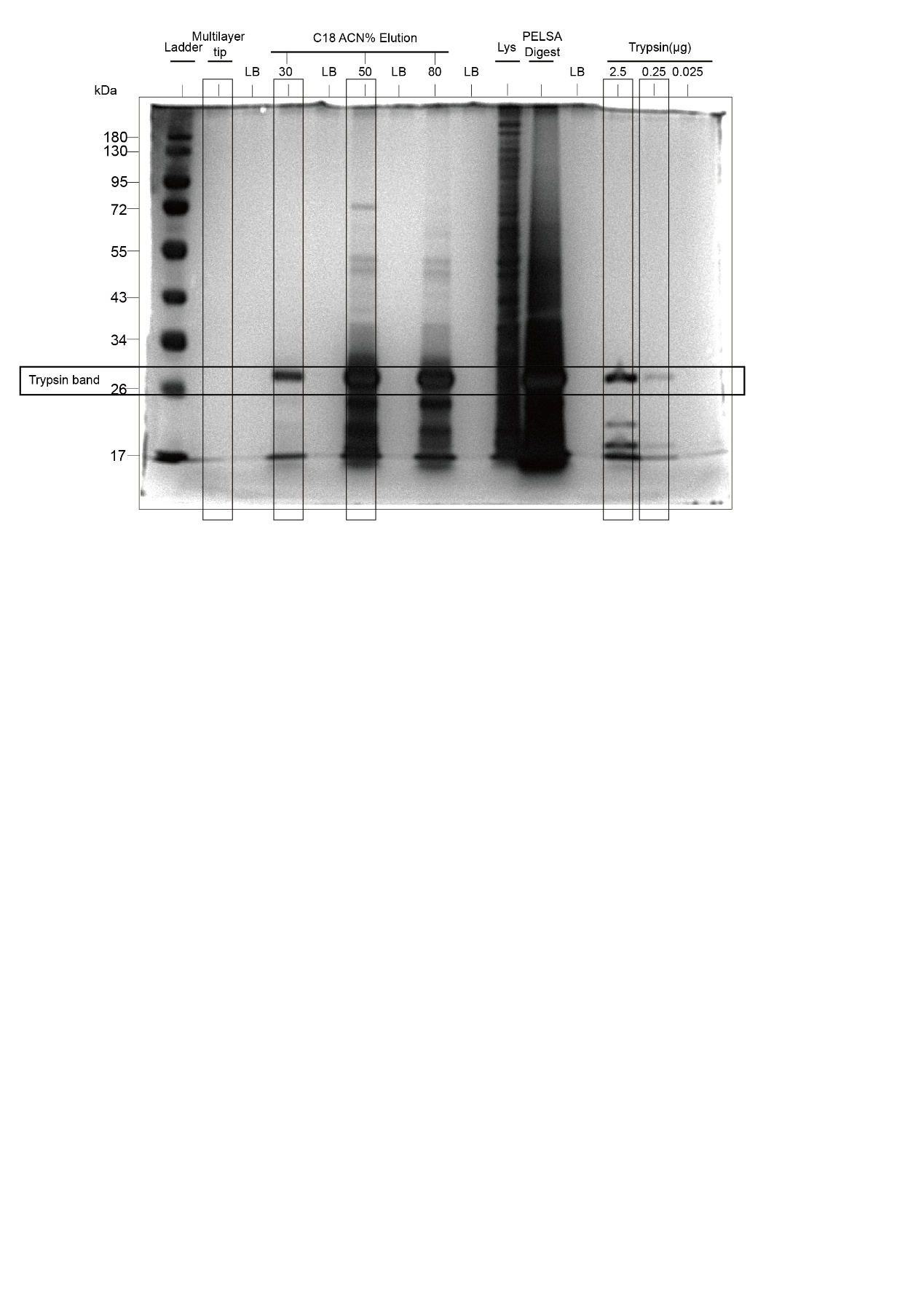

Figure S3. Optimization of cleanup steps in AutoPELSA. Silver-stained SDS–PAGE analysis comparing eluates generated from three distinct cleanup methods: the multilayer C4–SCX–C18 tip, a 5 mg C18 tip, and a 100 mg C18 SPE column. For the multilayer C4–SCX–C18 tip and the 5 mg C18 tip, the analyzed eluate corresponds to the 30% ACN fraction. For the 100 mg C18 SPE column, the analyzed eluate corresponds to the 50% ACN fraction. An additional 80% ACN fraction from the 5 mg C18 tip is also shown. The band corresponding to trypsin is clearly indicated. The definitions for the five bands indicated by black boxes, which appear in Figure 2b, are provided here. LB (Loading Buffer) represents the blank loading control. Lys represents the lysate prior to trypsin digestion. PELSA digest represents the lysate following trypsin digestion.

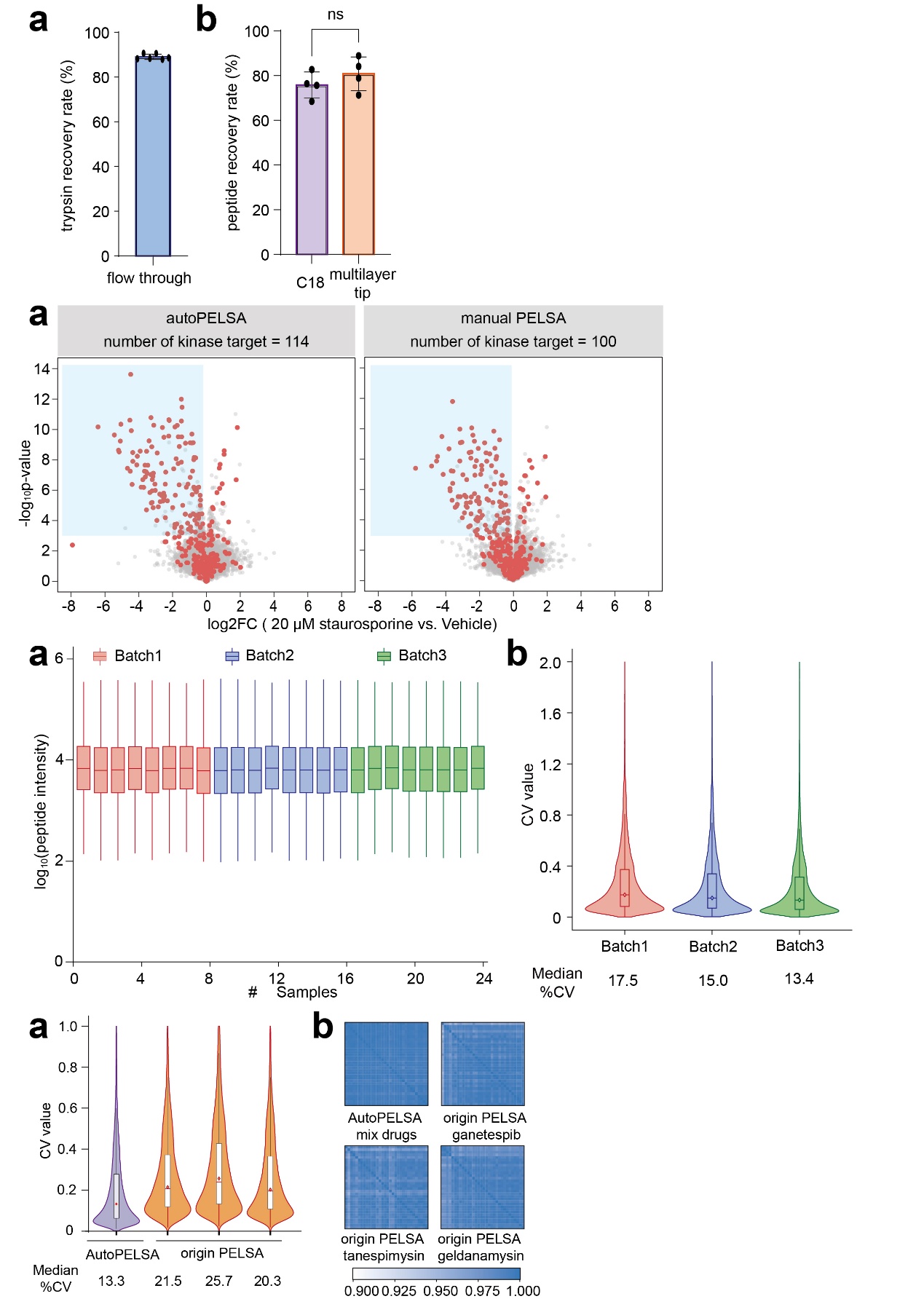

Figure S4. Proteome profiling performance using AutoPELSA. (a) Boxplots of log10-transformed peptide intensities across three batches. (b) Violin plots showing distributions of coefficients of variation (CVs) of peptide intensities across three batches.
